# Repeat expansion in a Fragile X model is independent of double strand break repair mediated by Pol θ, Rad52, Rad54l or Rad54b

**DOI:** 10.1101/2024.11.05.621911

**Authors:** Bruce E Hayward, Geum-Yi Kim, Carson J Miller, Cai McCann, Megan G. Lowery, Richard D. Wood, Karen Usdin

**Affiliations:** Section on Gene Structure and Disease Laboratory of Cell and Molecular Biology National Institute of Diabetes and Digestive and Kidney Diseases National Institutes of Health, Bethesda, MD 20892; The University of Texas MD Anderson Cancer Center Department of Epigenetics & Molecular Carcinogenesis PO Box 301429, Unit 1951, Houston, Texas 77230; Takeda Pharmaceuticals U.S.A., Inc. Global Biologics Informatics and Automation 500 Kendall Street, Cambridge, MA 02142, USA

**Keywords:** *FMR1*-related disorders (*FMR1* disorders), Fragile X-related disorders, Repeat Expansion Disease, Double-strand break repair, Gap-filling model

## Abstract

Microsatellite instability is responsible for the human Repeat Expansion Disorders. The mutation responsible differs from classical cancer-associated microsatellite instability (MSI) in that it requires the mismatch repair proteins that normally protect against MSI. LIG4, an enzyme essential for non-homologous end-joining (NHEJ), the major pathway for double-strand break repair (DSBR) in mammalian cells, protects against expansion in mouse models. Thus, NHEJ may compete with the expansion pathway for access to a common intermediate. This raises the possibility that expansion involves an NHEJ-independent form of DSBR. Pol θ, a polymerase involved in the theta-mediated end joining (TMEJ) DSBR pathway, has been proposed to play a role in repeat expansion. Here we examine the effect of the loss of Pol θ on expansion in FXD mouse embryonic stem cells (mESCs), along with the effects of mutations in *Rad52*, *Rad54l* and *Rad54b,* genes important for multiple DSBR pathways. None of these mutations significantly affected repeat expansion. These observations put major constraints on what pathways are likely to drive expansion. Together with our previous demonstration of the protective effect of nucleases like EXO1 and FAN1, and the importance of Pol β, they suggest a plausible model for late steps in the expansion process.

## Introduction

More than 45 human diseases have been shown to result from an expansion in the size of a disease-specific short tandem array or microsatellite ^1^. This group of diseases, known as the Repeat Expansion Diseases (REDs), includes the Fragile X-related disorders (FXDs) which result from the expansion of a CGG-repeat tract in the 5’ untranslated region of the X-linked *FMR1* gene ^2^. We have used a mouse model of the FXDs we had previously generated ^3^ to identify a number of mismatch repair factors that either protect against or promote repeat expansion ^4–10^. Many of these factors have also been shown to modify expansion in models of other Repeat Expansion Diseases (REDs) ^11–16^. Studies in several different REDs have identified some of these same genetic factors as modifiers of expansion risk in humans ^17–23^. This suggests that many of the REDs share the same expansion mechanism and that our FXD mouse model is suitable for studying this process.

Another clue to the expansion mechanism that has emerged from studies in the FXD mouse models is that LIG4, a ligase essential for non-homologous end-joining (NHEJ) ^24^, the major form of double-strand break (DSB) repair (DSBR) that operates in mammals, protects against expansion ^25^. A similar protective effect for LIG4 has recently been shown in an HD mouse model ^13^. In addition to NHEJ, LIG4 has recently been implicated in the restart of replication forks stalled by R-loops ^26^. However, since expansion can occur in both dividing and non-dividing cells via a process that is modulated by the same set of core proteins, it may be that LIG4’s protective effect reflects its role in NHEJ and thus that expansion involves a DSB intermediate or one that can be inter-converted with one.

Pol θ-mediated end-joining (TMEJ), the major form of alternative end-joining (Alt-EJ) used to repair DSBs in mammalian cells ^27,28^, has been proposed to play a role in the expansion process ^29,30^. Pol θ, an A-family DNA polymerase, has many properties that make it an appealing contributor to the expansion process. It is important for the repair of substrates with long 3’ overhangs ^27^ and TMEJ involves repair synthesis directed by short tracts of homology, typically shorter than those involved in single strand annealing (SSA) ^31^. Pol θ is often associated with templated insertions ^32,33^. Whilst most of these insertions are ∼3 bp in length, insertion tracts of 5-30 bp are commonly observed ^32,33^. Furthermore, Pol θ can tightly grasp a 3’ overhang through unique contacts in the active site, allowing the enzyme to extend substrates that have as few as 2-4 nt of annealed sequence ^32,33^.

RAD52, RAD54L and RAD54B are proteins that play vital roles in a variety of homology-driven repair processes. RAD52 is required for the repair of transcription-related DSBs ^34–36^, single stranded breaks ^37^ and for some forms of break-induced replication (BIR) ^38,39^, including the Mitotic DNA synthesis (MiDAS) occurring at common fragile sites ^40^. BIR has also been suggested to be responsible for repeat expansions seen in a mammalian tissue culture model system ^41,42^. RAD52 also plays an important role in the single-strand annealing (SSA) pathway ^43^ that involves the annealing of long regions of homologous repeat sequences that flank the break. A null mutation in *Rad52* was previously shown to have a minor effect on repeat expansion in a mouse model of myotonic dystrophy (DM1), another member of the REDs ^14^, but the role for this DSBR factor has not been examined in other model systems. Ablation of either *Rad54l* or *Rad54b* in mouse embryonic stem cells (mESCs) results either in a mild reduction in homologous recombination (HR) in the case of *Rad54l*, or no effect in the case of *Rad54b*. However, the absence of both RAD54L and RAD54B dramatically reduces the HR efficiency ^44^, consistent with the importance of these proteins in HR. A null mutation in *Rad54l* was found to have no effect on repeat expansion in the DM1 mouse model while the effect of a mutation in *Rad54b* or in both *Rad54l* and *Rad54b* was not examined ^14^.

Here we describe the effects of null mutations in *Polq*, *Rad52, Rad54l* and *Rad54b* on repeat expansion in a mESC model of the FXDs ^45^. We also studied lines with mutations in both *Rad54l* and *Rad54b*. We show that none of these DSBR factors modulate expansion in these cells. The exclusion of many homology-driven processes, provides insights into what are likely to be late steps in the expansion mechanism and allows to us to refine our model for repeat expansion.

## Materials and Methods

### Reagents

All standard reagents and tissue culture reagents were as previously described ^5^.

### Western blotting

Commercial antibodies for western blotting were as follows: RAD52, 1:1000 (ABclonal #A3077); RAD54, 1:200 (Santa Cruz, #sc-374598); ß-actin, 1:1000 (Abcam #ab8226); HRP-conjugated donkey anti-rabbit IgG, 1:2000 (GE Healthcare, #NA934); HRP-conjugated goat anti-mouse IgG, 1:10000 (Invitrogen #31430) / Supplemental Figure 1).

The QL fragment of Pol θ ^46^ was used to generate antibody 153-5-1 against Polθ. Purified Polθ antibody (2.4 mg/mL) was used at 1:500 dilution. This antibody (now available as #48160 from Cell Signaling Technologies) detects a >250 kDa protein that is absent in cells derived from *Polq^-/-^* mice (Supplemental Figure 1).

### CRISPR-Cas9 modification of FXD mESCs

The generation of mouse embryonic stem cells (mESCs) from a FXD mouse model was previously described ^45^. Mutations were generated in *Polq*, *Rad52*, *Rad54l* and *Rad54b* using either a dual guide RNA (gRNA) strategy for *Polq* or a single gRNA for the others as previously described ^5^ and illustrated in panel A of Figs 1-3. Edited lines were identified by PCR using distally located primers in adjacent introns or exons as indicated by the red arrows. Edits in lines that produced a PCR product were verified by cloning and sequencing of the PCR products. In cell lines where two edited alleles could not be sequence-verified the absence of the gene product was confirmed using western blotting whenever suitable antibodies were available. The repeat number in bulk DNA was determined from total genomic DNA isolated from ∼200,000 cells as previously described ^47^. CRISPR-edited lines and size-matched mock-edited cell lines were propagated together using previously described culture conditions ^5^ to ascertain their expansion rates. Small pool PCR was performed as previously described ^48^.

**Fig. 1.**
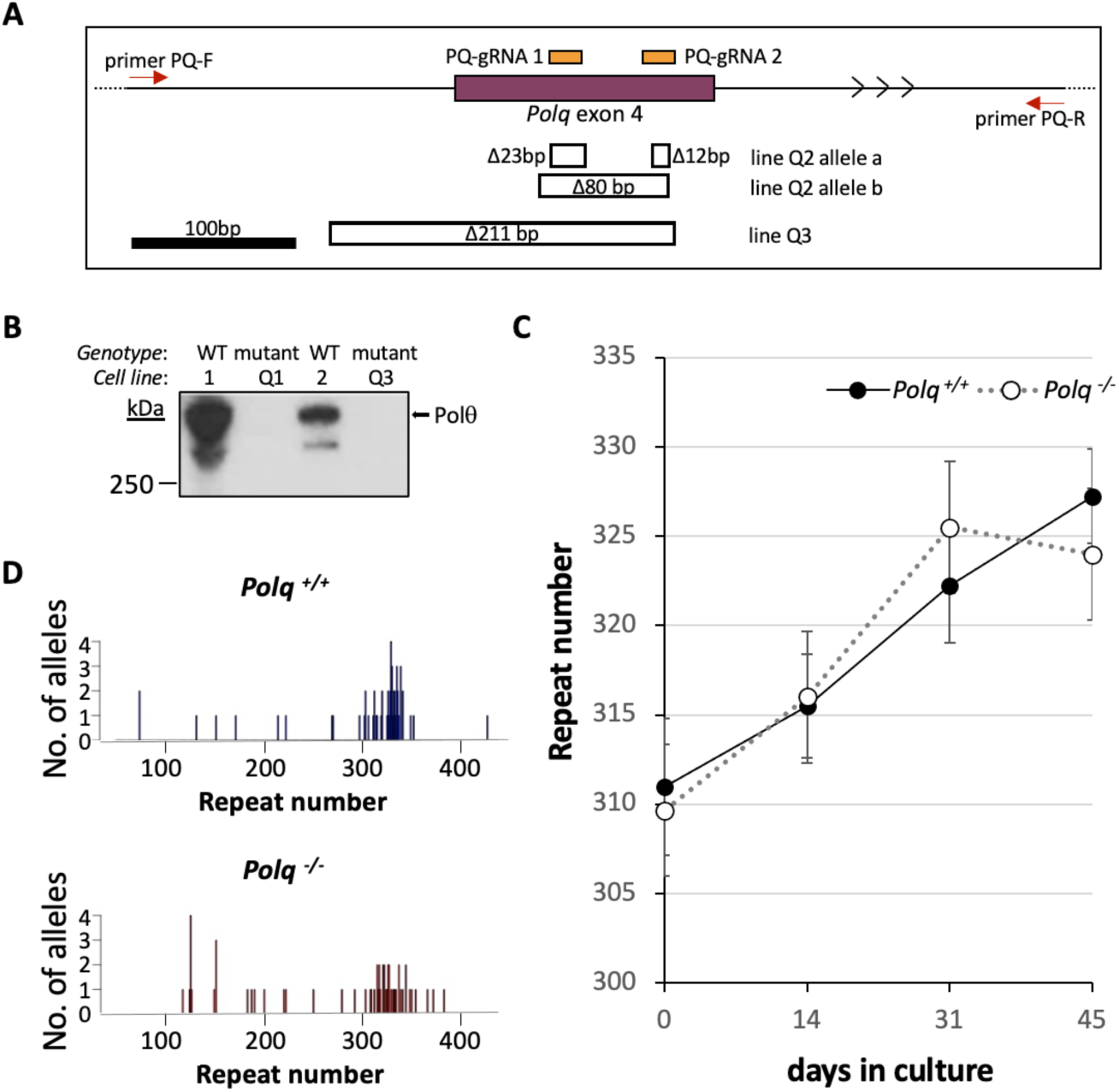
Effect of loss of Polθ on repeat expansion. A) Diagram of the region of *Polq* targeted for CRISPR mutation and the structure of the deletions at the target site. The positions of the two guide RNAs used to target exon 4 of *Polq* (chr16:37017199-37017355, mm10) are shown in orange. The location of the primers used for PCR to identify edited lines are shown by the red arrows. The extent of deletions identified in lines 17 and 22 are indicated by hollow boxes under the exon diagram. B) Western blot with Pol θ antibody of lines with alleles that could not verified by sequencing. C) Chart showing the average repeats added over time for size and passage matched *Polq* ^+/+^ and *Polq* ^-/-^ lines. The error bars represent the standard error. D) Small pool PCR of *Polq*^+/+^ and *Polq*^-/-^ lines after 44 days in culture.

**Fig. 2.**
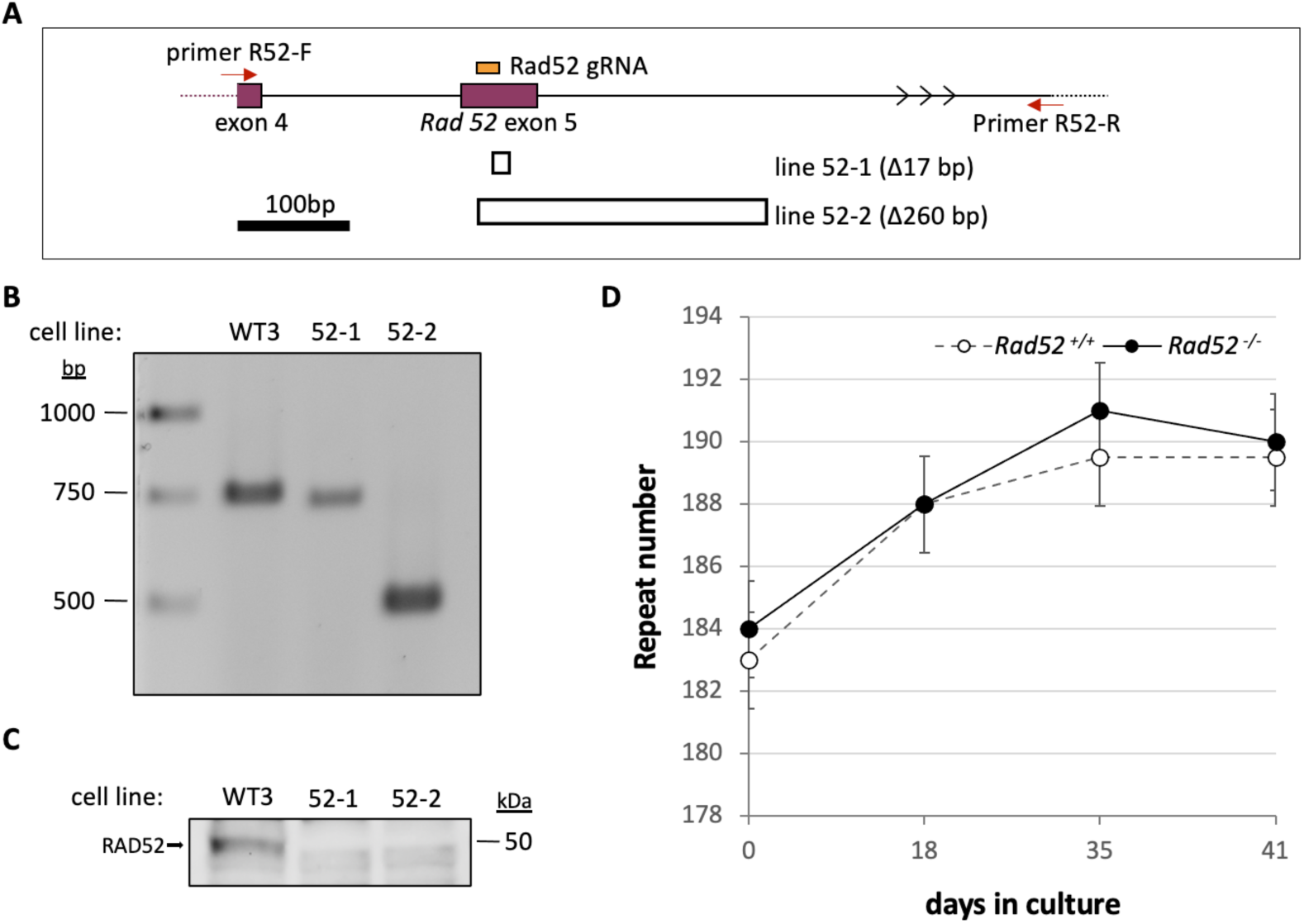
Effect of the loss of RAD52 on repeat expansion. A) Diagram of the region of *Rad52* targeted for CRISPR mutation and the structure of the deletions at the target site. The position of the gRNA used to target exon 5 of *Rad52* (chr6:119914170-119914237, mm10) is shown in orange. The locations of the primers used for PCR to identify edited lines are shown by the red arrows. The extent of deletions identified in lines C8 and C11 are indicated by hollow boxes under the exon diagram. B) Agarose gel of PCR products amplified from the indicated cell lines using primers 1 and 2, indicated above. C) Western blot of a *Rad52*^+/+^ line and the two *Rad52*^-/-^ lines using an anti-RAD52 antibody. D) Graph showing the average number of repeats added over time in culture for size- and passaged-matched *Rad52*^+/+^ and *Rad52*^-/-^ lines. The error bars represent the standard error.

**Fig. 3.**
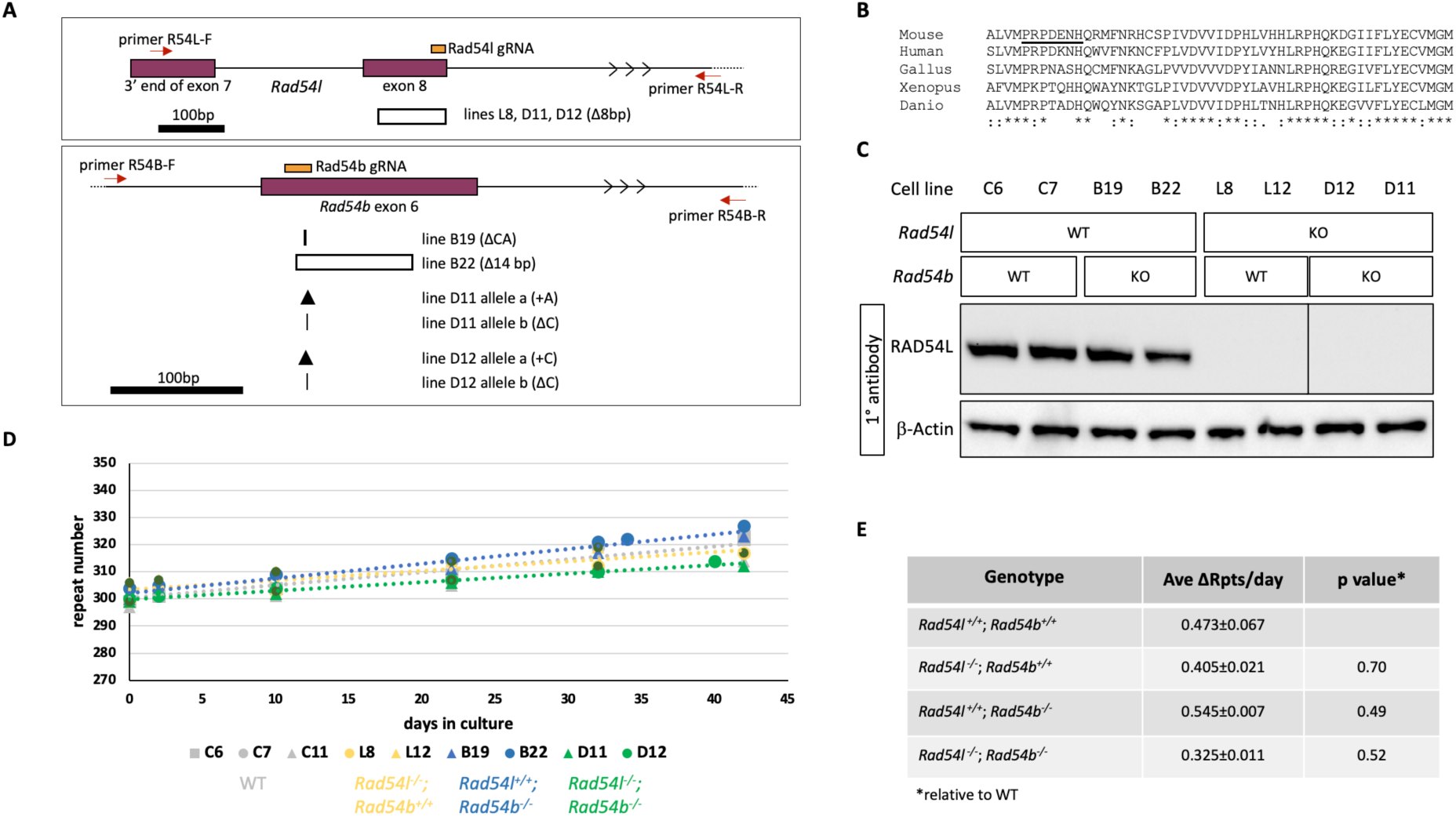
Effect of the loss of RAD54L and RAD54B on repeat expansion. A) Diagram of the regions of *Rad54l* and *Rad54b* targeted for CRISPR mutation and the structure of the deletions at the target sites. The positions of the gRNAs used to target exon 8 of *Rad54l* (chr4:116109969-116110093, mm10) and exon 6 of *Rad54b* (chr4:11597832-11597994, mm10) are shown in orange. The locations of the primers used for PCR to identify edited lines are shown by the red arrows. The extent of larger deletions is indicated by a hollow box. Smaller deletions are indicated by the vertical lines with the thicker line reflecting the loss of two bp and the thinner line the loss of a single bp. The arrowheads indicate the location of single base insertions. B) Alignment of the amino acids encoded by *Rad54b* exon 6 in various vertebrate species. * indicates sequence identity;: a conservative substitution and. a semi-conservative substitution. C) Western blot probed with anti-RAD54 antibody using β-actin as a loading control. KO, cell line treated with CRISPR reagents for the indicated gene. WT, cell line not exposed to CRISPR reagents for the indicated gene. Lysates were probed with the indicated antibodies. All samples probed with RAD54L antibody were analyzed on the same blot, but an unrelated sample loaded between lines L12 and D12 was cropped out. The unrelated sample was excluded from the β-actin-probed blot which was performed separately. *Rad54l* CRISPant cell lines show no detectable RAD54L protein and are presumed to be null mutants. D) Graph showing the change in repeat number over time in culture for size- and passaged-matched *Rad54l*^+/+^; *Rad54b*^+/+^ (lines C6, C7, C11), *Rad54l*^-/-^; *Rad54b*^+/+^ (lines L8, L12), *Rad54l*^+/+^; *Rad54b*^-/-^ (lines B19, B22) and *Rad54l*^-/-^; *Rad54b*^-/-^ (lines D11, D12) lines. E) Table showing the average repeats added over time in culture for size and passage matched WT and mutant lines. The p values were calculated as described in Methods, relative to the *Rad54l^+/+^; Rad54b^+/+^* control lines. Ave ΔRpts/day, number of repeats added per day averaged over all lines for the indicated genotypes, ± standard error.

### Statistics

The CGGC Permutation Test as implemented in the compareGrowthCurves function from the R package statmod was used to test for differences in the expansion rates between pairs of cell lines ^49,50^.

## Results

To examine the role of Pol θ in expansion we used a CRISPR-Cas9 dual guide RNA strategy to make deletions in exon 4 of the *Polq* gene in expansion-proficient mESCs derived from our FXD mouse model ^5,45,47^. We identified mutant cell lines by PCR using screening primers situated in intron 3 and intron 4 as illustrated in Fig. 1A. We focused on those lines with a similar repeat number that would be large enough to see expansions and for which size-matched controls were available. Three lines each with ∼300 repeats were chosen for further study: Q1, Q2, and Q3. Lines Q2 and Q3 produced a PCR product with primers flanking the gRNAs; Line Q1 did not, suggesting a large deletion encompassing one or both screening primer sequences. PCR products were cloned and individual clones sequenced. As shown in Fig. 1A, Line Q2 had two frame shift alleles: one with two small deletions and another with a single 80bp deletion. Only a single allele encoding a frame shift mutation was detected for Line Q3. Because our sequencing could not account for both *Polq* alleles in Lines Q1 and Q3, we confirmed the absence of Pol θ protein by western blotting (Fig. 1B). We then monitored the change in the allele size in these cells over time in culture using the size matched unmodified lines as controls. As can be seen in Fig. 1C, the *Polq^+/+^* lines cell line gained an average of 0.375 repeats/day and the *Polq^-/-^* lines an average of 0.35 repeats/day. This difference was not statistically significant (p=0.9).

Given Pol θ’s ability to generate larger templated insertions, we wondered if Pol θ was important for generating the small number of larger expansions that are seen in *Polq^+/+^* cells. Since these events are relatively rare, they are not apparent when the allele distribution is examined in bulk culture. To address this possibility, we carried out small pool PCR comparing the expansion products produced in *Polq^+/+^* and *Polq^-/-^* lines. The allele profiles are shown in Fig. 1D. In the *Polq*^+/+^ population 15% had gained 10 repeats or more, compared to 18% of the *Polq*^+/+^ population. This difference was not significant (Fisher’s exact test, p=0.79). There was also no significant difference in the distribution of the alleles either when the whole population was compared (Mann-Whitney, p=0.1031) or the expansions alone (Mann-Whitney, p=0.0872). This suggests that the loss of Pol θ does not significantly affect the production of large expansions in these lines.

To evaluate contributions to expansion from other NHEJ and TMEJ-independent DSB pathways, we first made *Rad52* null mutations in cell lines with ∼150 CGG-repeats using a single gRNA that targets exon 5. Edited lines were verified by PCR using a primer in exon 4 and a primer located in intron 5 as illustrated in Fig. 2A-B, followed by cloning and sequencing the PCR product. Line 52-1 showed a 17 bp deletion in one allele with no PCR product corresponding to the second allele. Line 52-2 had a single allele with a 260 bp deletion that removed the exon 5/intron 5 boundary. The absence of RAD52 in both lines was confirmed by western blot (Fig. 2C). Mutant cell lines were then examined over time in culture alongside size-matched unmodified cell lines. As can be seen in Fig. 2D, cell lines lacking RAD52 showed no statistically significant difference (p=0.83) in the rate of expansion relative to the unmodified cells with *Rad52*^+/+^ cells gaining ∼0.11 repeats/day and *Rad52*^-/-^ cells gaining ∼0.14 repeats/day.

To evaluate a role for RAD54L and/or RAD54B in either promoting or protecting against expansion, we mutated these genes using CRISPR-Cas9 and a single guide RNA in either exon 8 or exon 6, respectively, as illustrated in Fig. 3A. Exon 8 of *Rad54l* encodes a DExH/DEGH-box helicase domain important for RAD54 function and the gRNA is coincident with a sequence encoding highly conserved residues required for ATP binding. Exon 6 of *Rad54b* is also a highly conserved region that has 52% of its amino acids being invariant from zebrafish to man (Fig. 3B). We identified *Rad54l* edited cell lines by PCR using a primer in exon 7 and a second primer in intron 8 as illustrated in Fig. 3A. PCR products were then sequenced. One line, L12, had a large deletion on both alleles since no PCR product was obtained. A second cell line L8 had one allele with an 8 bp deletion. Western blotting confirmed that neither cell line produced RAD54L (Fig. 3C). In the case of the *Rad54b* edited lines, one line, B19, had a single detectable allele that had a 2 bp deletion. A second line, B22, had a single allele detected by PCR that had a 14bp deletion. There was no evidence for a wild-type *Rad54b* allele in either line. Unfortunately, no suitable antibodies were available to verify the loss of RAD54B protein. However, the absence of a PCR product from the second allele in each line would mean that the entire highly conserved exon 6 was lost (Fig. 3B). Furthermore, splicing of exons 5 and 7 would result in an in-frame stop codon. Thus, we conclude these alleles are likely to be functionally null. We also generated *Rad54l*^-/-^; *Rad54b*^-/-^ lines by using CRISPR-Cas9 to make *Rad54b* null mutations in the *Rad54l*^-/-^ line L8 using the gRNA used to generate the *Rad54b^-/-^* lines. Two cell lines were identified, D11 and D12, each with two frameshifted alleles in *Rad54b*: one with a single base added and one with a single base deleted.

We then monitored these lines for change in repeat number over time in tissue culture together with unmodified cell lines with comparable repeat numbers. As can be seen in Fig. 3D and E, the singly-mutated cell lines expand at similar rates to the unmodified control lines. The average expansion rate of the doubly mutant lines is slightly lower than the unedited or single mutant lines, but this difference does not reach statistical significance for any pairwise comparison between the double mutant lines and other lines (p values ranged from 0.21 to 0.83). Thus, it is apparent that none of the proteins tested are essential for expansion. The small effect of the knockout of both *Rad54l* and *Rad54b* may reflect a subtle contribution of these proteins to non-HR pathways or perhaps even the indirect effect of HR proteins on gene expression ^51,52^.

## Discussion

We have shown that the loss of Pol θ (Fig. 1), RAD52 (Fig. 2), or the loss of either or both RAD54L and RAD54B (Fig. 3) had no significant effect on the extent of expansion in the mESC model of FXD repeat expansion. The lack of an effect of the loss of Pol θ suggests that TMEJ does not play an essential role in either promoting or preventing expansion. Furthermore, the fact that the loss of RAD52 or the loss of both RAD54L and RAD54B have little or no effect on the repeat expansion rate in FXD mESCs suggests that many other homology-dependent DSBR pathways-including HR, SSA and synthesis-dependent strand annealing-are also not required either for expansion in this model system or for protection against it. The lack of an effect of the loss of RAD52 would also exclude expansion models invoking RAD52-dependent BIR ^41^. Furthermore, it would also suggest that RAD52-dependent repair of transcription-related DSBs ^34–36^ does not explain the dependence of expansion on transcription ^10^. The lack of a role for many homology-dependent DSBR processes is consistent with our previous demonstration that homozygosity for the *Mre11*^ALDT1/ALDT1^ mutation does not affect the expansion frequency ^25^ and that 5’ to 3’ exonucleases like EXO1 and FAN1 are protective ^6,7^. Taken together it suggests that DSBR mechanisms requiring extensive end-resection are not responsible for expansions in this model system.

One way to reconcile the observation that LIG4 is protective is if at some frequency the expansion process generates an intermediate that can be repaired either by NHEJ or via a form of Alt-EJ that does not require Pol θ. Pol β has recently been shown to participate in a form of Pol θ-independent Alt-EJ that competes very effectively with NHEJ for repair of DSBs ^53^. A role for Pol β-mediated Alt-EJ is appealing as a source of expansions since *in vitro* experiments show that this polymerase can efficiently repair relatively short 3’ overhangs containing other trinucleotide repeats in such a way as to generate expansions ^54^. Furthermore, we have shown that Pol β is an important contributor to expansions in our FXD mouse model ^55^.

We speculate that following MutLγ cleavage of an expansion substrate consisting of loop-outs formed on both strands of the repeat (Fig. 4 (i)), the nicks might be processed as in normal MMR, either by 5’ to 3’ exonuclease resection followed by gap-filling by Pol θ; or in the absence of exonuclease processing, by Pol θ-mediated strand-displacement, flap removal and gap-filling (Fig. 4 (ii-iii)). We hypothesize that, at some frequency, excessive exonuclease processing of the nicks on each strand results in an intermediate like that shown in Fig. 4 (iv), in which the promoter proximal and promoter distal ends of the repeat can dissociate. The ends retain the potential to reanneal because of the sequence homology in each of the 5’ overhangs. Further end-processing could result in a substrate suitable for NHEJ (Fig. 4 (v)). Alternatively, reannealing of the two ends could occur with subsequent gap-filling by an enzyme like Pol β (Fig. 4 (vi)), that we have previously shown to promote expansions ^55^. In principle, the outcome of this form of Alt-EJ could be a contraction, an unchanged allele, or an expansion involving the gain of a variable number of repeats. The outcome would depend on the extent of the overhang and the size of any gaps that arise after reannealing. Since limited EXO1 processing of nicks would be predicted to generate expansions like those generated by Pol θ acting alone, a model such as this might provide an explanation for EXO1’s protective effect.

**Fig. 4.**
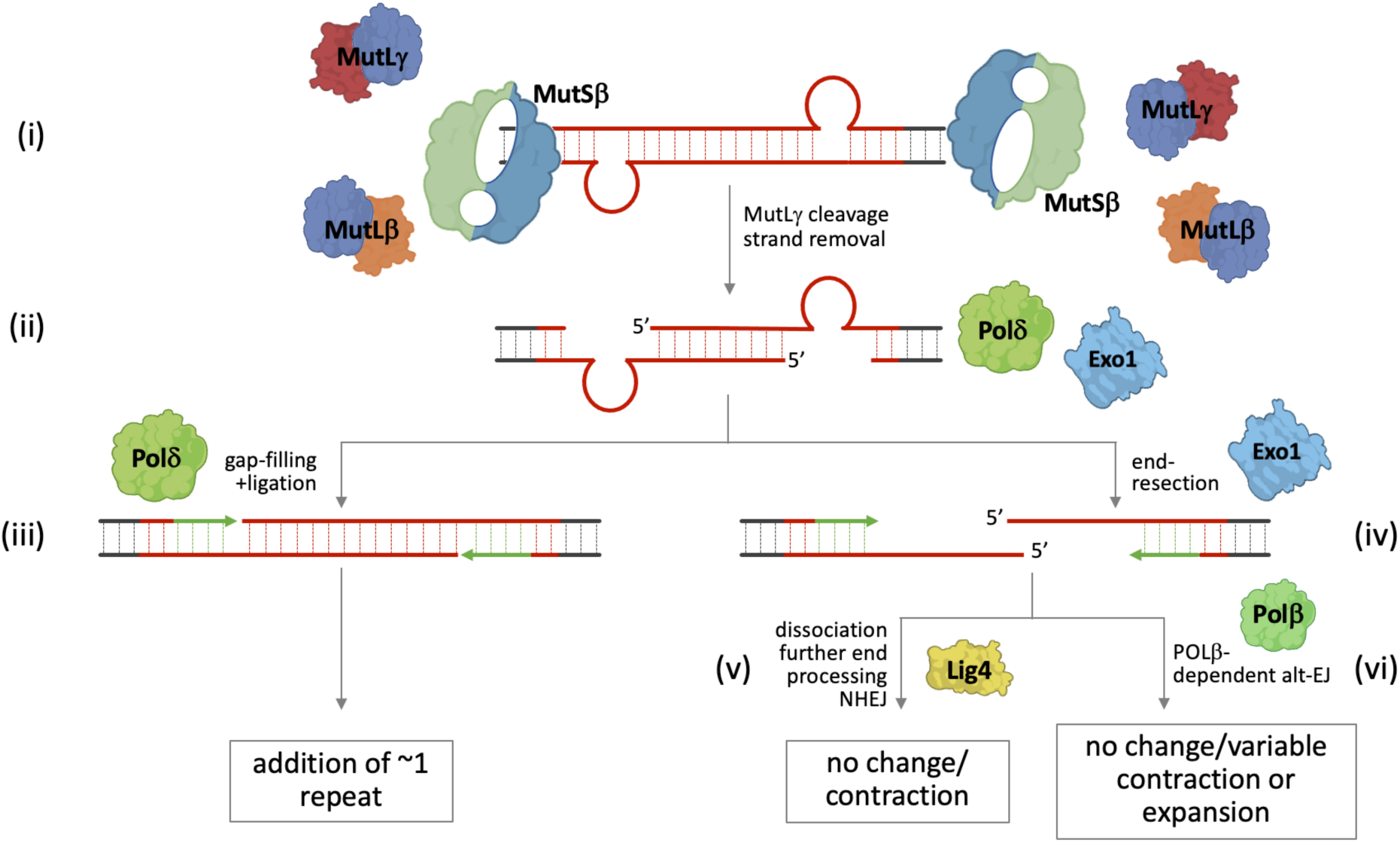
Model for repeat expansion. (i) Expansion is initiated by the formation of offset loop-outs on each strand perhaps as a result of out-of-register reannealing during transcription. (ii) Binding of MutSb, MutLb and MutLψ to each loop-out with subsequent MutLψ cleavage results in a nick opposite each loop-out that is processed either by a 5’ to 3’ exonuclease like EXO1 or by strand-displacement by Pol θ followed by flap removal. (iii) The resultant gap opposite each loop-out can be filled in by Pol θ to generate a small expansion. (iv) However, at some frequency, exonucleolytic processing is so extensive that the two ends of the repeat dissociate. Processing of the resultant 5’ overhangs can occur either by NHEJ resulting in contractions or unchanged alleles (v) or by reannealing and Pol β-mediated Alt-EJ to restore the original allele or result in a contraction or expansion with the change in repeat number depending on the size of the overhangs and any gaps present (vi).

## Acknowledgements

The authors would like to acknowledge the excellent bioinformatics support received from Dr Hernan Lorenzi of the NIDDK Trilabs Bioinformatics group.

## Author contributions

BH, G-YK, CJM, CM, MGL acquired and analyzed the data and helped draft the manuscript. RDW and KU, helped design the experiments, analyze the data and edit the manuscript.

---

**Table 1.**
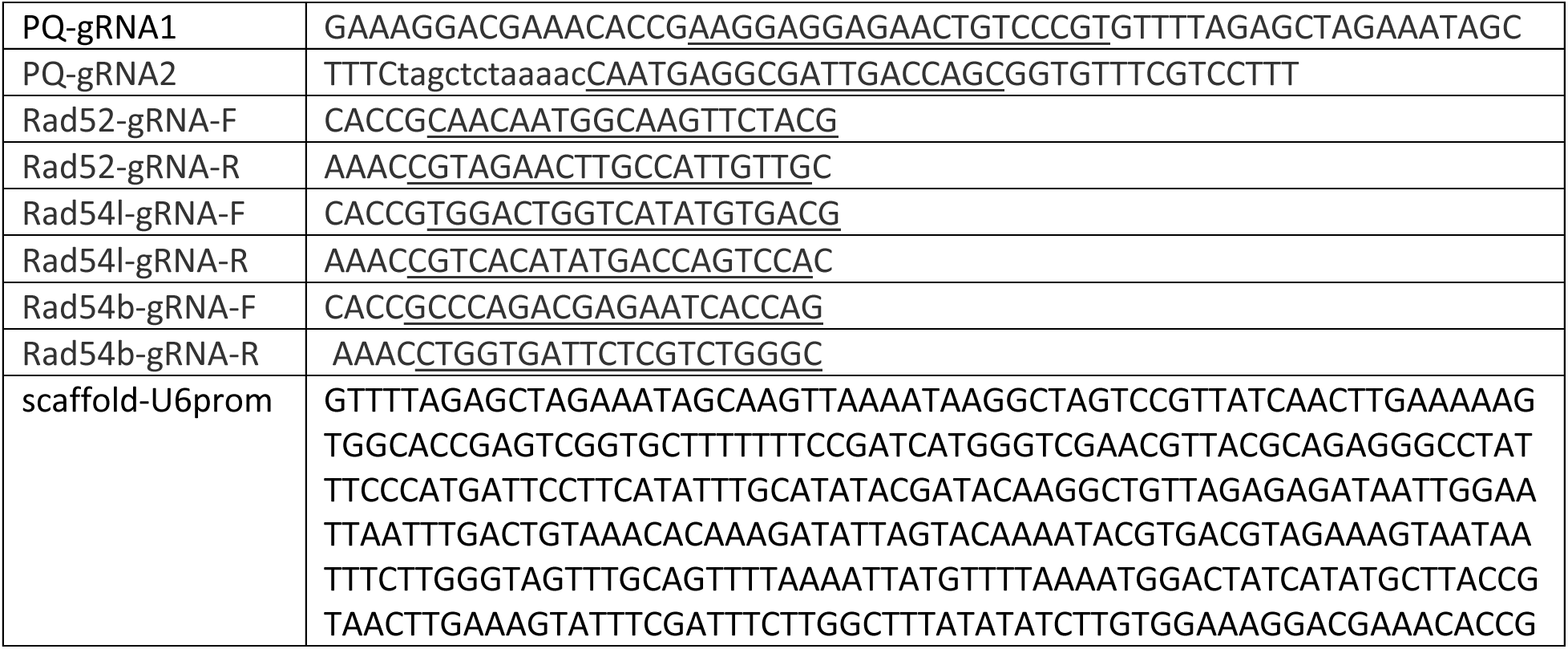
Primers and DNAs used for the CRISPR-Cas9 modifications. (gRNA sequences are underlined).

**Table 2.**
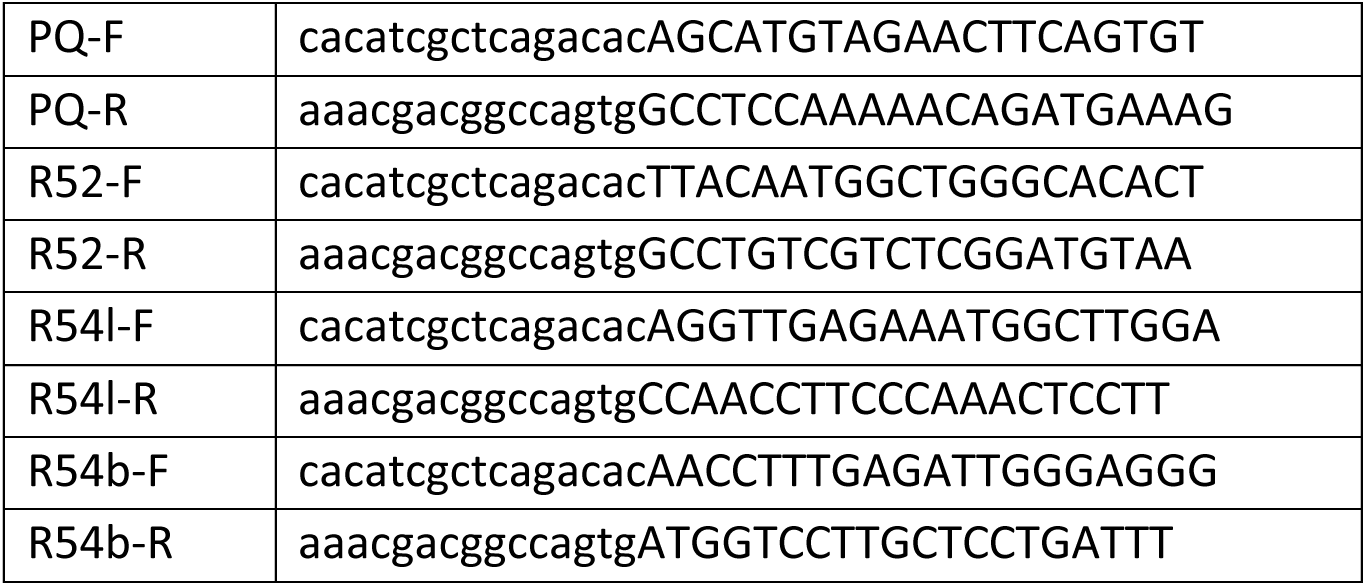
Primers used for screening CRISPR-targeted sites. (genome-specific regions capitalized).

**Supplemental Figure 1.**
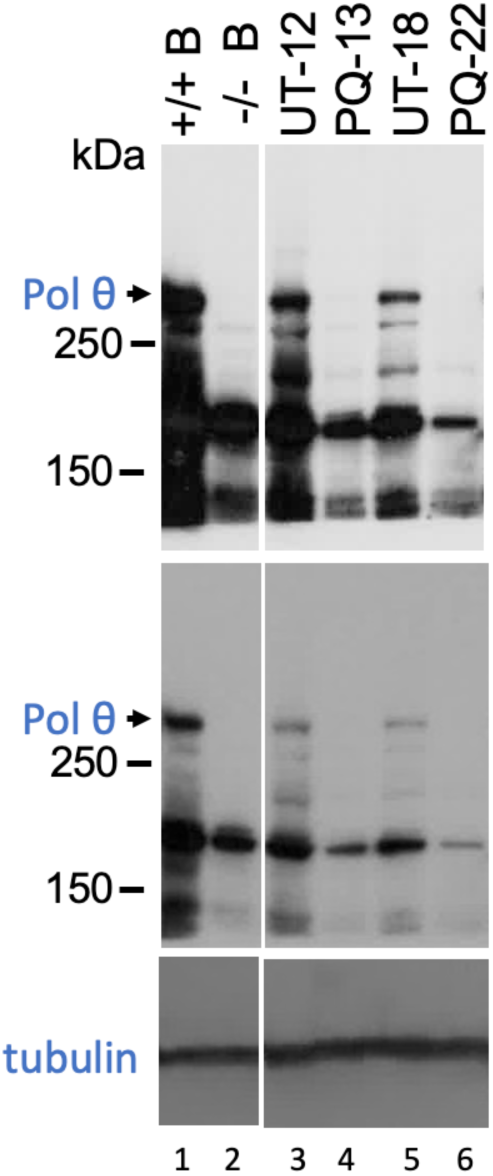
Confirmation of *Polq* gene knockout in mouse cell lines. An uncropped immunoblot of extracts from the indicated cell lines, using an antibody recognizing mouse Polθ. Extracts were prepared from B cells from *Polq*^+/+^ mice (lane 1); B cells from *Polq*^-/-^ mice (lane 2); *Polq*^+/+^ mouse cell line WT-12 (lane 3); *Polq*^-/-^ mouse cell line PQ-13 (lane 4); *Polq*^+/+^ mouse cell line WT-18 (lane 5); *Polq*^+/+^ mouse cell line PQ-22 (lane 6). The top two panels show darker and lighter exposures of the same immunoblot. Intervening lanes containing irrelevant samples were excised from the immunoblot as indicated. The 50 kDa region of the gel was cut away and immunoblotted separately with an antibody against alpha tubulin, shown in the bottom panel.

## Supplemental Methods

Extracts were prepared by lysis of pellets from 5 x 10^6^ cells. Cells were resuspended in buffer containing SDS, boiled for 10 min, sonicated, and insoluble material was removed by centrifugation. Samples (15 μL for B cells, 10 μL for other samples) were loaded on a 3-8% gradient Criterion™ XT Tris-acetate protein gel and run with XT running buffer at 75 V for 180 min. Markers were All Blue Precision Plus Protein standards (Bio-Rad). Following transfer and drying of the membrane, immunoblotting used mouse monoclonal antibody against Pol θ. This antibody (153-5-1) was raised against a fragment of Pol θ QL ^46^. Purified antibody (2.4 mg/mL) was used at 1:500 dilution in blocking buffer. Following washing, secondary goat anti-mouse HRP antibody was used at 1:10,000 dilution in 5% non-fat dried milk (CST) in TBS-T solution. The film was developed with Clarity plus reagent and exposed to x-ray film. A separate identical gel was loaded and stained with Revert total protein stain and imaged on a Li-Cor to confirm equal staining.

### Preparation of mouse B-cell extracts

*Polq*^-/-^ mice, originally derived by Shima *et al*. ^56^, were obtained from Jackson Laboratories as described and maintained on a C57BL/6J background ^57^. Naïve mouse B cells were isolated from spleens, negatively sorted with anti-CD43 beads, and cultured with lipopolysaccharide and interleukin-4 as described ^58^. Extracts were prepared after 72 h culture.

